# Speed modulation of hippocampal theta frequency and power predicts water maze learning

**DOI:** 10.1101/2020.03.31.016907

**Authors:** Calvin K. Young, Ming Ruan, Neil McNaughton

## Abstract

Theta oscillations in the hippocampus have many behavioural correlates, with the magnitude and vigour of ongoing movement being the most salient. Many consider correlates of locomotion with hippocampal theta to be a confound in delineating theta contributions to cognitive processes. But, theory and empirical experiments suggest theta-movement relationships are important if spatial navigation is to support higher cognitive processes. In the current study, we tested if variations in speed modulation of hippocampal theta can predict spatial learning rates in the water maze. Using multi-step regression, we find the magnitude and robustness of hippocampal theta frequency versus speed scaling can predict water maze learning rates. Using generalised linear models, we also demonstrate that speed and water maze learning are the best predictors of hippocampal theta frequency and power. Theta oscillations recorded from the supramammillary area showed much weaker, or non-existent, relationships, which supports the idea that hippocampal theta has specific roles in speed representation and spatial learning. Our findings suggest movement-speed correlations with hippocampal theta frequency may be actively used in spatial learning.

## Introduction

The functional significance of hippocampal theta (4-12 Hz) oscillations has remained an enigma since its initial characterisation (Green and Arduini, 1954). While the mechanisms of its generation are still incompletely understood (Bland, 1986; Buzsaki, 2002), the integrity of medial septal afferents (Brugge, 1965; Winson, 1978; Rawlins et al., 1979) appears to be crucial (but see Goutagny et al., 2009). Hippocampal theta has many behavioural correlates (Vanderwolf, 1969; Buzsaki, 2005; Young, 2011; Korotkova et al., 2018), the most salient among them is its linear scaling with the speed of ongoing locomotion, in both frequency (McFarland et al., 1975; Slawinska and Kasicki, 1998; Jeewajee et al., 2008; Wells et al., 2013; Aghajan et al., 2017) and power (Rivas et al., 1996; Maurer et al., 2005; Watrous et al., 2011; Ahmed and Mehta, 2012). Despite various replications in independent laboratories, linear locomotion speed and hippocampal theta correlation can be highly variable between animals and recording sessions (Richard et al., 2013; Hernandez-Perez et al., 2016). It has been suggested that variability in the speed-theta correlations may be modulated by other functions that are dependent on theta oscillations, such as learning and memory (Montgomery et al., 2009; Richard et al., 2013; Schmidt et al., 2013; Belchior et al., 2014; Hernandez-Perez et al., 2016). Certainly, abolishing hippocampal theta oscillations do not result in the abolishment of movement, but is correlated to spatial learning deficits (Winson, 1978; McNaughton et al., 2006). Electrically driving theta-paced stimulation in the hippocampus partially rescues hippocampal-dependent learning deficits (McNaughton et al., 2006; Shirvalkar et al., 2010). Computational models integrating spatial representations in the hippocampal-entorhinal circuit explicitly require speed modulation of theta oscillations in their contribution to path integration and spatial navigation (Burgess and O’Keefe, 2011), suggesting linear speed may be a partial correlate in its involvement in spatial navigation.

The linear theta-speed relationship can be degraded through uncoupling self-motion and actual translation through space, as indicated by studies using a running wheel (Czurko et al., 1999; Shin and Talnov, 2001), treadmills (Kuo et al., 2011), passive movement through carts (Terrazas et al., 2005) and virtual reality (Ravassard et al., 2013). Although swimming is natural and a normal mode of locomotion in rodents, past studies have indicated a lack of theta-speed relationship in swimming rats (Whishaw and Vanderwolf, 1973), presumably due to a disruption of proprioception associated with conventional ambulation on land. A more detailed study on hippocampal theta frequency and swimming speed have revealed an unreliable relationship at best (Hernandez-Perez et al., 2016) – although the use of conventional FFT-based spectral analysis in this study may have yielded less accurate estimates due to the non-stationary nature of brain field potentials and the time-frequency trade-off (Young and Eggermont, 2009).

Given the theoretical (Burgess and O’Keefe, 2011) and empirical (Brandon et al., 2011; Koenig et al., 2011; Bolding et al., 2019) support for hippocampal theta to contribute to spatial representation and spatial learning (McNaughton et al., 2006), we sought to carry out a more in-depth analysis on how hippocampal theta oscillations are related to locomotion/swimming parameters (i.e. speed and acceleration) in the water maze, and how the relationship may be linked to learning. We show that hippocampal theta frequency and power, but not those recorded from the supramammillary area, are differentially scaled with speed but less so with acceleration. The reliability and the magnitude of hippocampal theta versus swimming speed scaling can predict water maze learning rates. Together, our data support the notion that hippocampal theta-speed/acceleration relationships provide information relevant to path integration in water maze learning.

## Materials and methods

### Subjects

Eight male Sprague Dawley rats were acquired from the Department of Laboratory Animal Sciences at the University of Otago. These eight rats were taken from a larger dataset for a different experiment (McNaughton et al., 2006; Ruan et al., 2017), and were kept in a temperature and humidity controlled room with 12 hr light/dark cycles (lights on at 6 a.m.). All rats had water and food access *ad libitum* throughout the experiment. The rats were given a ten-day acclimatisation period before surgical procedures. Relevant to the current study, stainless steel bipolar recording electrodes (70 μm) were implanted in the hippocampus (AP: −3.8 mm, ML: −2.5 mm, DV: −3.5 mm with tip separation of 1 mm) and the supramammillary area (AP: −4.8 mm, ML: −.9 mm, DV: −9.4 mm with tip separation of .5 mm and implanted @ 6° from vertical) under ketamine/ medetomidine (75 mg/kg and .5 mg/kg, respectively) anaesthesia. Six anchor screws, one of which acted as the ground with uninsulated silver wire (.25 mm), held the implant with dental cement. A guide cannula and a stimulation electrode aimed at the medial septum and the fornix, respectively, were also implanted (McNaughton et al., 2006) but were not used in rats reported in the current study. Rats were given atipamezole (2.5 mg/kg) and postoperative analgesia at the end of the surgery and allowed to recover for at least ten days before any further manipulations. All procedures reported here were approved by the University of Otago Animal Ethics Committee (84/00 and 67/03).

### Water maze learning

We implemented a one-day water maze training protocol (Pan and McNaughton, 1997). A circular black pool (diameter: 150 cm; height: 35 cm) was filled with 26° (± 2°C) water to 25 cm deep. A black square 15 cm platform was placed 1.5 cm beneath the water and has a fixed position in the centre of the nominal “southeast” quadrant. The rats were connected to the recording system (see below) and allowed 40 s to find the platform. If successful, the rats remained on the platform for a further 15 s. If unsuccessful, the rats were guided to the platform. In either scenario, the next trial started immediately after 15 s on the platform. A counterbalance sequence of (NSWE ENSW WENS SWEN) 16 trials were administered. Position tracking and water maze performance were carried out in HVS Image (HVS Image, UK).

### Neurophysiological recordings, spectral analysis and statistics

Local field potentials (LFPs) were acquired through a custom-made unity gain headstage and amplified (Grass P511K at 1-30 Hz band-pass). The data were digitised (100 Hz) by a Micro1401 (CED, UK) with Spike2 software. Spectral analysis was performed using the Chronux package (Bokil et al., 2010) with 5 tapers and a numerical bandwidth of 3. All data analyses were carried out in Matlab (Mathworks, Natick, MA). Statistics are reported as mean ± standard error unless otherwise stated. Raw *p*-values are reported for all statistical tests unless *p*-values are <.001, where they are reported as *p* < .001. A false discovery rate was calculated at a threshold of *p* = .04.

### Instantaneous frequency and power

Raw LFP data were first z-scored prior to further processing. For frequency and amplitude assessments, we opted to use analytic signal derivatives for instantaneous frequency and power for near real-time estimation of these variables. HPC and SuM LFPs were first band-passed (5-12 Hz, zero-phase) with an acausal filter and then Hilbert transformed. The phase of the signals was smoothed by a 3^rd^ order Savitzky-Golay filter and its derivative is taken as the frequency estimate. The absolute value of the signal was taken as the envelope/power of the signal. Since there was a ten-fold difference in acquisition rate between neurophysiological and tracking data, both instantaneous frequency and envelope of the signals were binned in 10 samples and the median value in the bins was taken as the final estimates corresponding to a single tracking coordinate. The binned instantaneous frequency and envelope are referred to as *frequency* and *power*; collectively as *theta variables* throughout this report.

### Position, speed, and acceleration

Position data were extracted from the text output from HVS Image (HVS Image, UK). To minimise the misrepresentation of position, speed, and acceleration, we employed a 3-point median filter to correct minor tracking errors. However, we left extended large jumps between points, no tracking data, and clear poor tracking data as “no data” to avoid introducing bias to “instantaneous” position, speed and acceleration estimates. Two trials from the dataset were missing too much data and since they appeared to contain extensive mis-tracks by visual inspection were discarded. Tracking data from one trial was missing and hence the trial was also omitted. In total, 125 trials from eight rats were included in the final analyses. Distance, speed, and acceleration were all calculated based on the known pixel-to-centimetre relationship from the tracking coordinates.

In this study, we elected to use the conventional binned approach to examine the relationship between theta frequency/power and speed/acceleration (Wells et al., 2013; Young and McNaughton, 2020). Although rats essentially swam continuously inside the water maze, we decided to apply a low cut-off of 2.5 cm/s as in previous studies to: 1) facilitate direct comparisons to speed-theta relationships for land locomotion and; 2) to exclude periods where rats vigorously attempt to climb up the maze wall without spatial translation immediately after being placed into the pool during early stages of learning. In addition, we excluded the first 2 s of data from each trial to eliminate tracking errors relating to the experimenter placing rats into the pool, as the tracking was triggered by any high-contrast entity entering the camera’s field of view. In sum, we employed 2.5 cm/s speed bins across 5 to 45 cm/s and 0.5 cm/s/s acceleration bins across −3 to 3 cm/s/s for speed and acceleration estimates. These are collectively termed *kinematic* variables throughout the current study.

### Spatial distribution and cross-correlations

Spatial distributions of kinematic and theta variables were constructed by first converting the value found at each coordinate into a “map” for each trial, where repeated samples of the same variable at the same coordinates represented by the average found in the spatial bin. Each raw spatial bin approximated a 20 x 20 cm area in the pool. All matrices from all trials were averaged to form a single map for each variable, convolved with a 3 x 3 gaussian filter and up-sampled 3 fold to yield a final map for 2D cross-correlation analysis.

### Correlating speed/acceleration theta modulations and learning

For linking theta variables to kinematic variables, we took the averaged theta variables across all available trials from each rat as primary data. Linear regression was applied to each individual rat’s data, as well as at the population level across rats. Slope and correlation coefficients were extracted from each fit.

Learning was quantified by extracting the exponential fit coefficient over the path length for each rat over the 16 trials. The slope and correlation coefficients of the theta frequency versus speed regressions from each individual rat were then regressed against their respective exponential fit coefficient of water maze performance over time. We used these series of regressions to demonstrate the robustness and reliability of theta frequency versus speed relationship with water maze learning.

### Instantaneous estimate of water maze performance

To provide an “instantaneous” estimate of water maze performance, we introduced a performance metric taking into account the direction of the rats’ trajectory, distance to the platform and normalised elapsed time in the trial. Particularly, we took the angle differences between each pair of tracking points to estimate the change in trajectory angle (*θ*_*point*_), and the angle differences between each tracking point to the point representing the centre of the platform (*θ*_*platform*_). The absolute difference between these two angles (i.e. point-to-point and point-to-platform) yielded a value between 0 and π for headings towards the platform and 180° away, respectively. This angle variable is then multiplied by the distance from the point to the platform (*D*_*platform*_) and added to the time remaining till the end of the trial (by locating or being guided to the platform; *T*_*end*_):

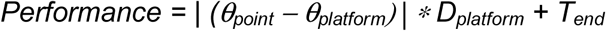

This metric favours correct head trajectories towards the platform over the absolute distance from the platform to account for varied starting points (i.e. distances from the hidden platform) for each trial. The addition of time remaining in the trial offsets spurious trajectory angles toward the hidden platform during the search.

### Generalised linear model

To gauge the individual and combined contribution of swimming kinematics and trajectory (i.e. performance) to theta measures, we entered z-scored binned swimming speed, binned acceleration and the performance metric as predictor variables so such:

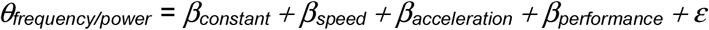

We extracted the fit parameters (i.e. slopes; standardised β, or zβ from hereon) for each rat and tested if the population averages are significantly different from zero with Student’s t-tests.

### Histology

At the end of the experiments, rats were euthanised with a lethal dose of sodium pentobarbital and transcardially perfused with physiological saline followed by 10% formalin in saline solution. After a minimum of 24 hr 10% formalin fixation, the brains were dehydrated with 30% sucrose in 10% formalin solution until saturation and sectioned at 90 μm on a freezing microtome. Electrode tracks were reconstructed from digitised Nissl stained sections.

## Results

### Continuous hippocampal and supramammillary theta in the water maze

We were able to confirm the placement of bipolar electrodes to have predominantly targeted the CA1 region of the hippocampus (Figure 1A, left panel) and the supramammillary area (Figure 1A, right). Both areas showed strong, continuous theta rhythmicity during swimming in the water maze by visual inspection (Figure 1B, left panels) and spectral analyses (Figure 1B, right panels). The presence of high amplitude, continuous theta oscillations throughout swimming circumvents the possibility our analytic signal-derived theta variables contain erroneous estimates of non-theta states.

**Figure 1.**
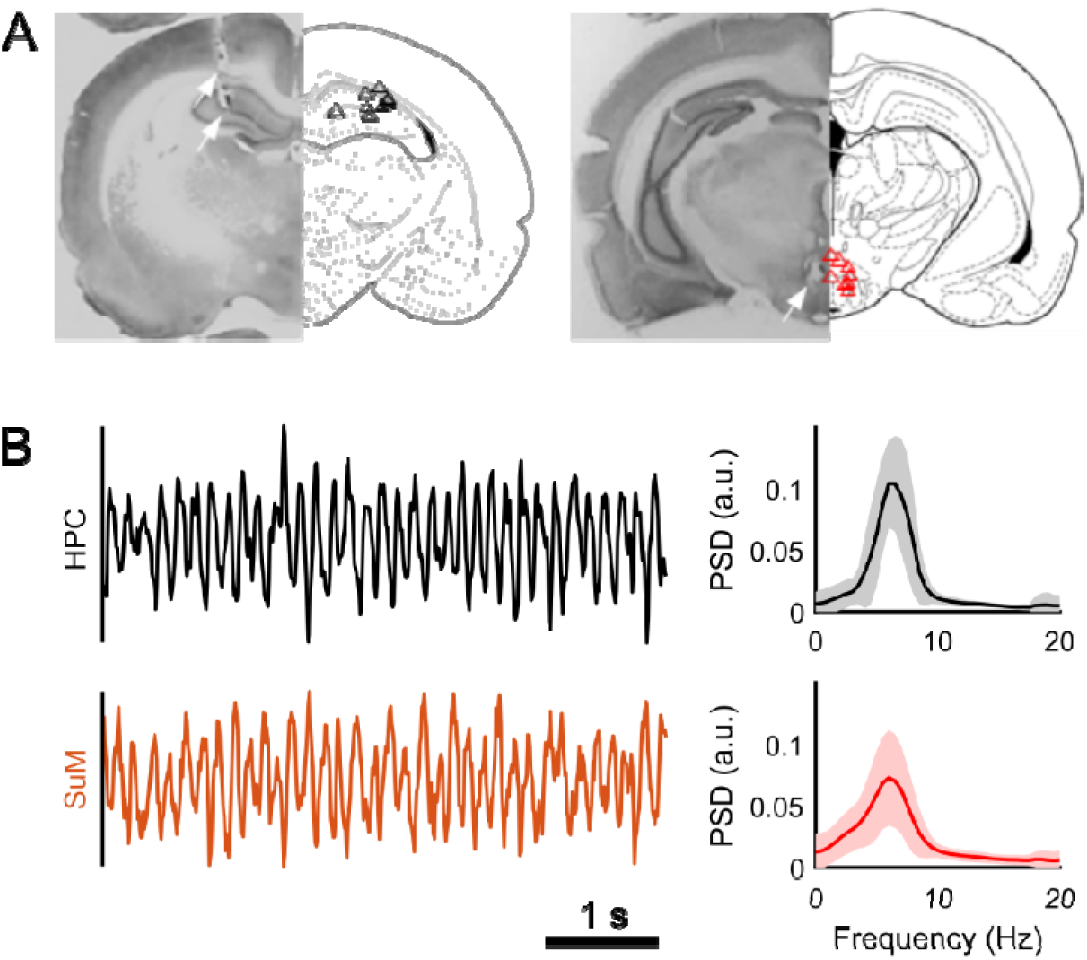
Theta field potential recordings. **A**: Example histology and reconstructed recording locations in the hippocampus (left) and the supramammillary area. **B**: Field potentials in the hippocampus (top, black) and the supramammillary area (bottom, red) show continuous theta oscillations during swimming.

### Spatial distribution of locomotion and theta parameters

To examine how theta oscillations relate to behaviour in the water maze, we first mapped theta variables (i.e. frequency and power) and kinematic measures (i.e. speed and acceleration) in the pool. In general, there do not appear to be any differences between HPC and SuM theta frequency distributions at the periphery but HPC theta frequency may be lower (Figure 2A, left) in the inner annulus while SuM theta frequency appears to be higher (Figure 2A, right). For theta power, the HPC seems to have higher power in general (Figure 2B, left) compared to SuM (Figure 2B, right). Speed (Figure 2C, left) distribution indicates low speeds in the target quadrant, particularly around the platform itself, which is consistent with qualitative observations that rats generally slow down near the hidden platform as part of the searching strategy. The speed distribution also suggests that the fastest speeds are observed in the “western” half of the pool. Acceleration (Figure 2C, right) shows a similar distribution to speed in that the highest changes are away from the periphery and on the “western” side of the pool. The platform is surrounded by a ring of deceleration, which is surrounded by a larger ring of acceleration, again corroborating qualitative observations of rats swimming quickly towards the platform and slow down as they approach the platform as a search strategy. As expected, the periphery of the pool had the highest occupancy (Figure 2D) as rats spent most of their actual time in the water initially in thigmotaxis. There is also a clear increase in occupancy around the hidden platform (but not “south-east” from it), again consistent with observed finer searching behaviour in the vicinity of the hidden platform once rats acquire its location.

**Figure 2.**
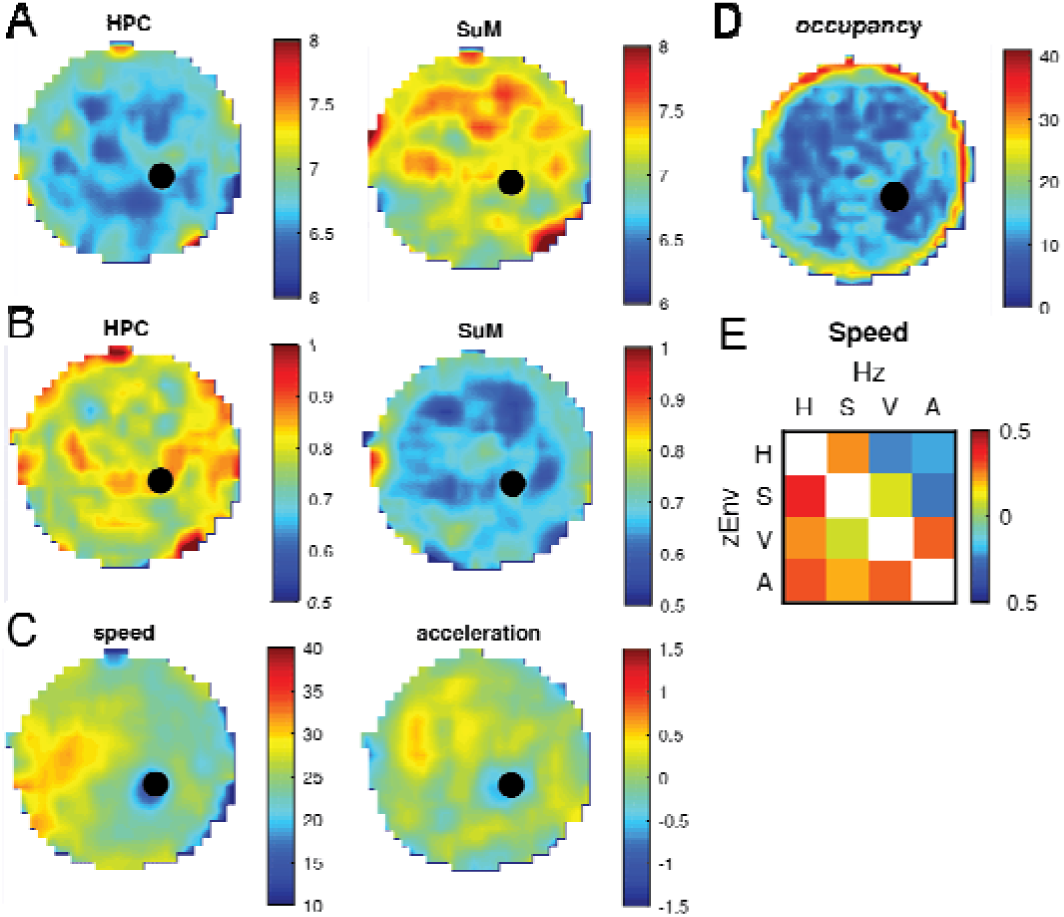
Spatial distribution of theta and kinematic variables in the water maze. **A**: Averaged theta frequency distribution within the water maze in the hippocampus (left) and the supramammillary area (right). **B**: Same as in **A** but for normalised power. **C**: Averaged swimming speed (left) and acceleration (right) distributions within the water maze. **D**: Occupancy map in the water maze. **E**: Spatial cross-correlation matrix with frequency (Hz) based cross-correlations on the top left half of the matrix and power (zEnv) based cross-correlations on the bottom right half. H: hippocampus; M: supramammillary area; V: speed; A: acceleration. Color maps *r*^*2*^ from −0.5 to 0.5.

While some general patterns of theta and swimming kinematic parameters can be qualitatively described, we performed spatial autocorrelation to see if additional features exist. Apart from the already described faster speeds from the “western” side of the pool towards the middle and high occupancy around the periphery, no other spatial autocorrelation features emerged (data not shown). We further explored potential similarities in spatial distribution across all theta and kinematic parameters in the pool using spatial cross-correlation (Figure 2E). There is a pattern of negative spatial cross-correlation between HPC theta frequency with swimming speed (*r* = .21) and acceleration (*r* = .22). There is weak positive spatial correspondence between HPC and SuM theta measures (*r* = .21 for frequency; *r* = .38 for power), as well between speed and acceleration (*r* = .28). There is minimal correlation between SuM theta power with speed (*r* = .1) or acceleration (*r* = .02), while SuM theta frequency showed no spatial correspondence with speed (*r* = .09) but a negative correlation with acceleration (*r* = −.26) and HPC theta power distribution showed positive spatial correspondence with speed (*r* = .24) and weak negative correlation with acceleration (*r* = −.13).

### Water maze learning

Before examining the relationship of theta variables with swimming kinematics and water maze learning, we first characterise metrics for learning. The decrease of path length as training progressed was evident for all but one rat (Figure 3A). An exponential decay function was fit for all “learning curves” (Figure 3B) and the slope parameter was extracted as a measure for water maze learning rate in analyses described in the next sections. We also introduced a moment-to-moment index of water maze performance as a metric for learning (see Methods). In Figure 3C, an example metric for the whole 40 s at the beginning of the training session (i.e. trial 1) is depicted, with its corresponding trajectory mapped in Figure 3E. In the last trial, where rats exhibited much shorter path length to reach the hidden platform, the metric accounts for erroneous trajectories away from the hidden platform and regress towards zero (Figure 3D) as rats successfully locate the platform (Figure 3F).

**Figure 3.**
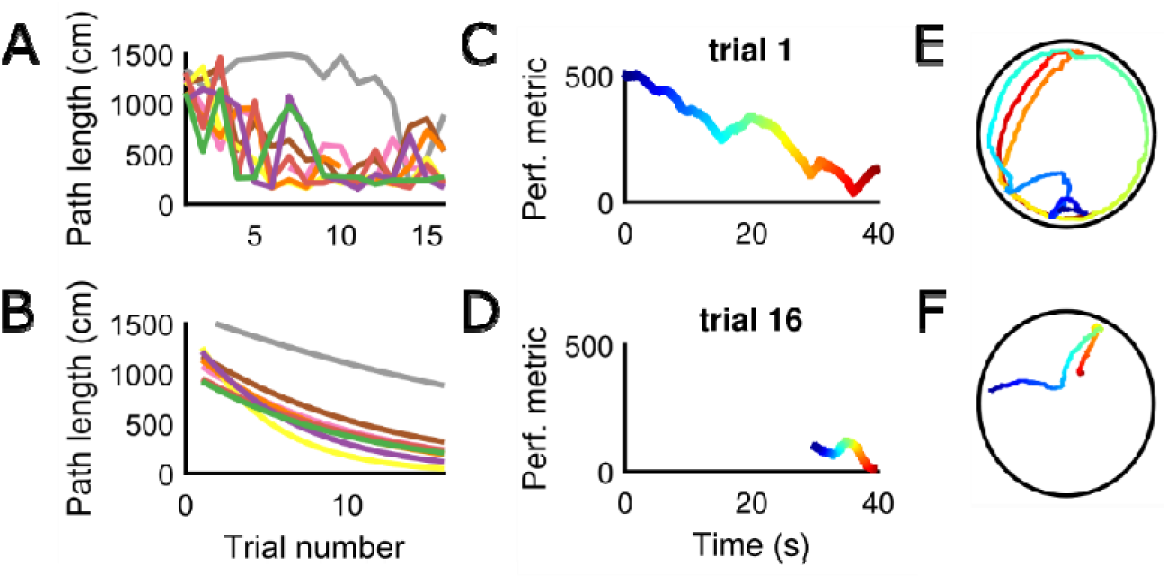
Characterisation of learning metrics in the water maze. **A**: Change of path length across trials for all rats. **B**: Associated exponential fits for **A. C**: An example of the water maze performance metric for the first exposure to the water maze. **D**: An example of the water maze performance metric in the last trial. **E**: Actual swim path in the water maze with color mapping to **C. F**: Same as in **E** but for **D**.

### Frequency and power correlates to speed

In the water maze, all rats except for one showed a nominal positive correlation between swimming speed and HPC theta frequency (Figure 4A, left). More variable regression lines for SuM theta frequency versus swimming speed (Figure 4A, middle) suggest a lack of relationship between the two variables. At the population level, HPC theta frequency and swimming speed do share positive linear correlation (*r*^*2*^ = .42, F(15) = 10.98, *p* = .005) while a more modest negative correlation was observed for SuM theta frequency versus swimming speed (*r*^*2*^ = .30, F(15) = 6.57, *p* = .022). To explore if better linear scaling between theta frequency and swimming speed correlates to learning, we regressed the slope (Figure 4B, top panels) and *r*^*2*^ (Figure 4B, bottom panels) of individual theta frequency versus swimming speed linear fits against learning (i.e. slopes of exponential fits from Figure 3B). Both the slope (Figure 4B, top left; *r*^*2*^ = .54, F(6) = 7.11, *p* = .037) and *r*^*2*^ (Figure 4B, bottom left; *r*^*2*^ = .71, F(6) = 15.02, *p* = .008) showed better learning associated with stronger/more reliable theta frequency versus swimming speed scaling. A similar but much weaker trend can be seen for the SuM equivalents for both slope (Figure 4B, top right; *r*^*2*^ = .22, F(6) = 1.67, *p* = .236) and *r*^*2*^ (Figure 4B, top left; *r*^*2*^ = .43, F(6) = 4.48, *p* = .079)

**Figure 4.**
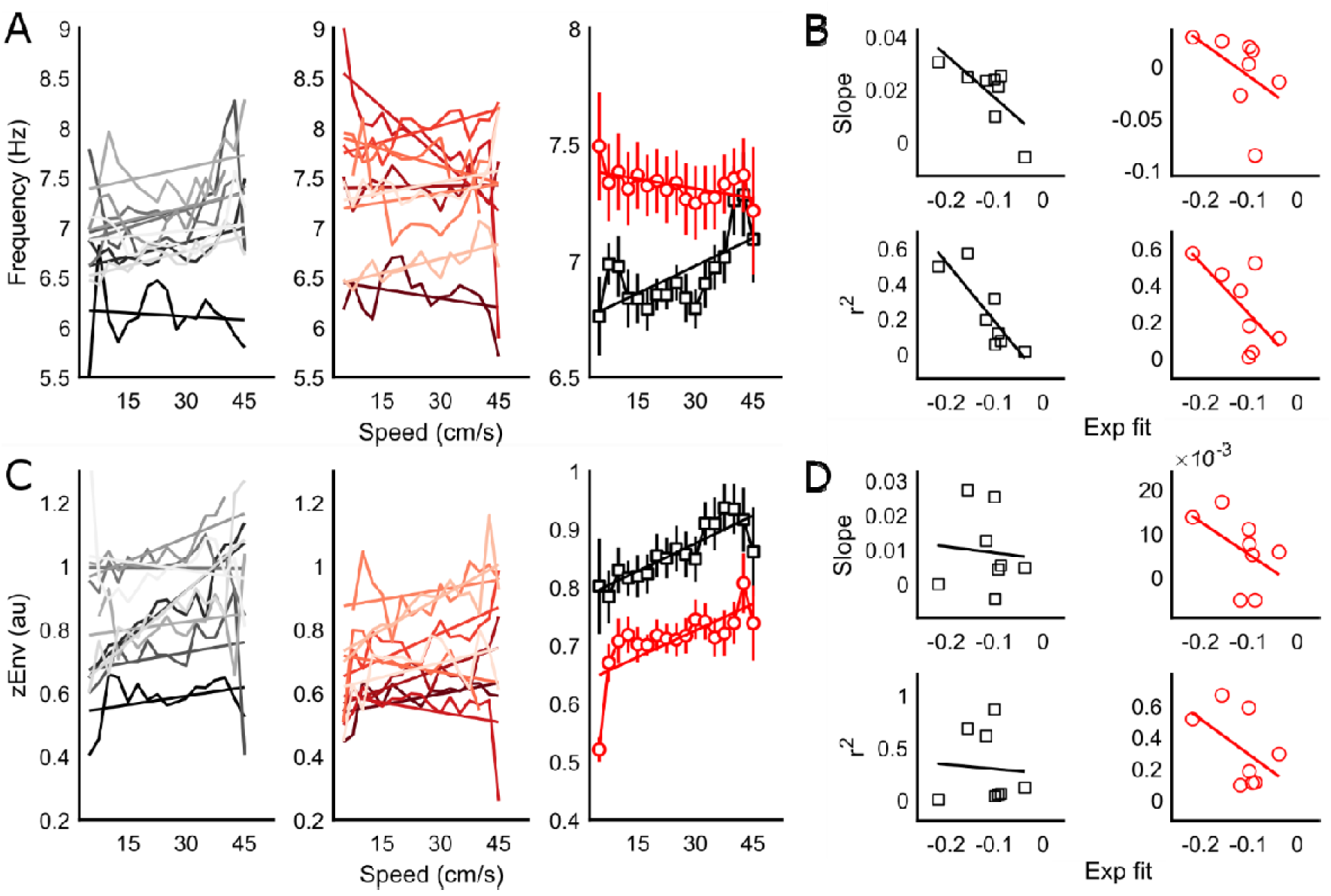
Speed modulation of hippocampal and supramammillary theta oscillations. **A**: Raw data and their linear fits for the hippocampus (right, black) and the supramammillary area (middle, red) theta frequency across binned speeds. Population data and their fits are depicted in the right panel. **B**. Correlations between the magnitude (slope, top) and robustness (*r*^*2*^, bottom) of the frequency versus speed scaling and water maze learning (exponential fit slope parameter) for hippocampal (black, left) and supramammillary (red, right) theta. **C**: Same as in **A** but for speed versus theta power correlations. **D**: Same as in **B** but for theta power versus speed scaling and water maze performance correlations.

The theta power versus swimming speed relationships in the HPC across all rats were consistently positive, except for one rat (Figure 4C, left). As with frequency, theta power versus swimming speed also yielded high variability in individual fits but mostly positive (Figure 4C, middle). At the population level, both HPC (*r*^*2*^ = .74, F(15) = 43.44, *p* < .001) and SuM (*r*^*2*^ = .47, F(15) = 13.23, *p* = .002) theta power showed a positive correlation with swimming speed (Figure 4C, right). Unlike with theta frequency, neither slope (*r*^*2*^ = .007, F(6) = .04, *p* = .85) nor *r*^*2*^ (*r*^*2*^ = .004, F(6) = .02, *p* = .88) of theta power versus swimming speed linear regressions from the HPC correlated with water maze learning (Figure 4D, left panels). Slope (*r*^*2*^ = .23, F(6) = 1.83, *p* = .22) and *r*^*2*^ (*r*^*2*^ = .26, F(6) = 2.06, *p* = .20) for SuM theta power versus swimming speed had better linear fits than their HPC counterparts, but were not statistically significant (Figure 4D, right panels). A summary between linear scaling of theta variables and their relationship with water maze learning are presented in Table 1.

**Table 1.** Summary from Figures 4B, 4D, 5B and 5D. *r*^*2*^ with *p* < .05 are highlighted in bold. Hz: theta frequency; zEnv: theta power.

| Hz | HPC |  | SuM |  |
| --- | --- | --- | --- | --- |
| | slope | $r^2$ | slope | $r^2$ |
| speed | **0.5425** | **0.7146** | 0.218 | 0.4276 |
| accel. | 0.0127 | 0.369 | **0.6545** | **0.5607** |

| zEnv | HPC |  | SuM |  |
| --- | --- | --- | --- | --- |
| | slope | $r^2$ | slope | $r^2$ |
| speed | 0.0069 | 0.0041 | 0.234 | 0.2556 |
| accel. | 0.0477 | 0 | 0.0907 | 0.0204 |

### Frequency and power correlates to acceleration

Individual linear fits for HPC theta frequency versus acceleration yielded mostly negative slopes with a single exception (Figure 5A, left). The slopes of linear fits in the SuM is also predominantly negative with more variability (Figure 5B, middle). Previous studies have shown both HPC theta frequency (Shin and Talnov, 2001) and power (Long et al., 2014) may show better positive correlation with deceleration than acceleration. Given the shallow slopes at the individual level linear regression, we opted to perform population level theta measure versus acceleration linear regressions separately for positive acceleration and negative acceleration, in addition to a single fit encompassing acceleration and deceleration. A summary of linear scaling of theta variables by negative, positive and overall acceleration is presented in Table 2. Overall linear regression of HPC theta frequency versus acceleration (Figure 5A, left) reveal a negative correlation (*r*^*2*^ = .68, F(10) = 23.09, *p* < .001), with negative acceleration (*r*^*2*^ = .79, F(10) = 19.04, *p* = .007) presumably contributing more to the negative slope than positive acceleration (*r*^*2*^ = .02, F(10) = .09, *p* = .78). In the SuM, the overall fit has a marginal negative slope (*r*^*2*^ = .24, F(10) = 3.55, *p* = .09) with theta frequency weakly correlating to the magnitude of negative (*r*^*2*^ = .19, F(10) = 1.19, *p* = .33) and positive (*r*^*2*^ = .47, F(10) = 3.56, *p* = .13) acceleration. Since the overall fit provided the best fit compared to negative or positive acceleration fits in both the HPC and the SuM, we regressed the individual slope (Figure 5B, top left; *r*^*2*^ = .01, F(10) = .08, *p* = .79) and *r*^*2*^ (Figure 5B, bottom left; *r*^*2*^ = .37, F(10) = 3.51, *p* = .11) from the overall fits to water maze learning (i.e. slope of the exponential fit over path length over time) and found no relationship. SuM theta frequency versus acceleration fit slopes (Figure 5B, top right; *r*^*2*^ = .65, F(10) = 11.36, *p* = .02) and *r*^*2*^ (Figure 5B, bottom right; *r*^*2*^ = .56, F(10) = 7.66, *p* = .04) correlated with water maze learning.

**Table 2.** Summary of overall, positive (+ve) and negative (-ve) acceleration fits with theta variables relating to Fig. 5. *r*^*2*^ with *p* < .05 are highlighted in bold. Hz: theta frequency; zEnv: theta power.

| Hz | HPC | SuM | zEnv | HPC | SuM |
| --- | --- | --- | --- | --- | --- |
| | $r^2$ | $r^2$ | | $r^2$ | $r^2$ |
| overall | **0.6773** | 0.2442 | overall | 0.0085 | 0.0573 |
| +ve | 0.0211 | 0.471 | +ve | **0.9765** | 0.3958 |
| -ve | **0.792** | 0.1917 | -ve | **0.6163** | **0.8544** |

**Figure 5.**
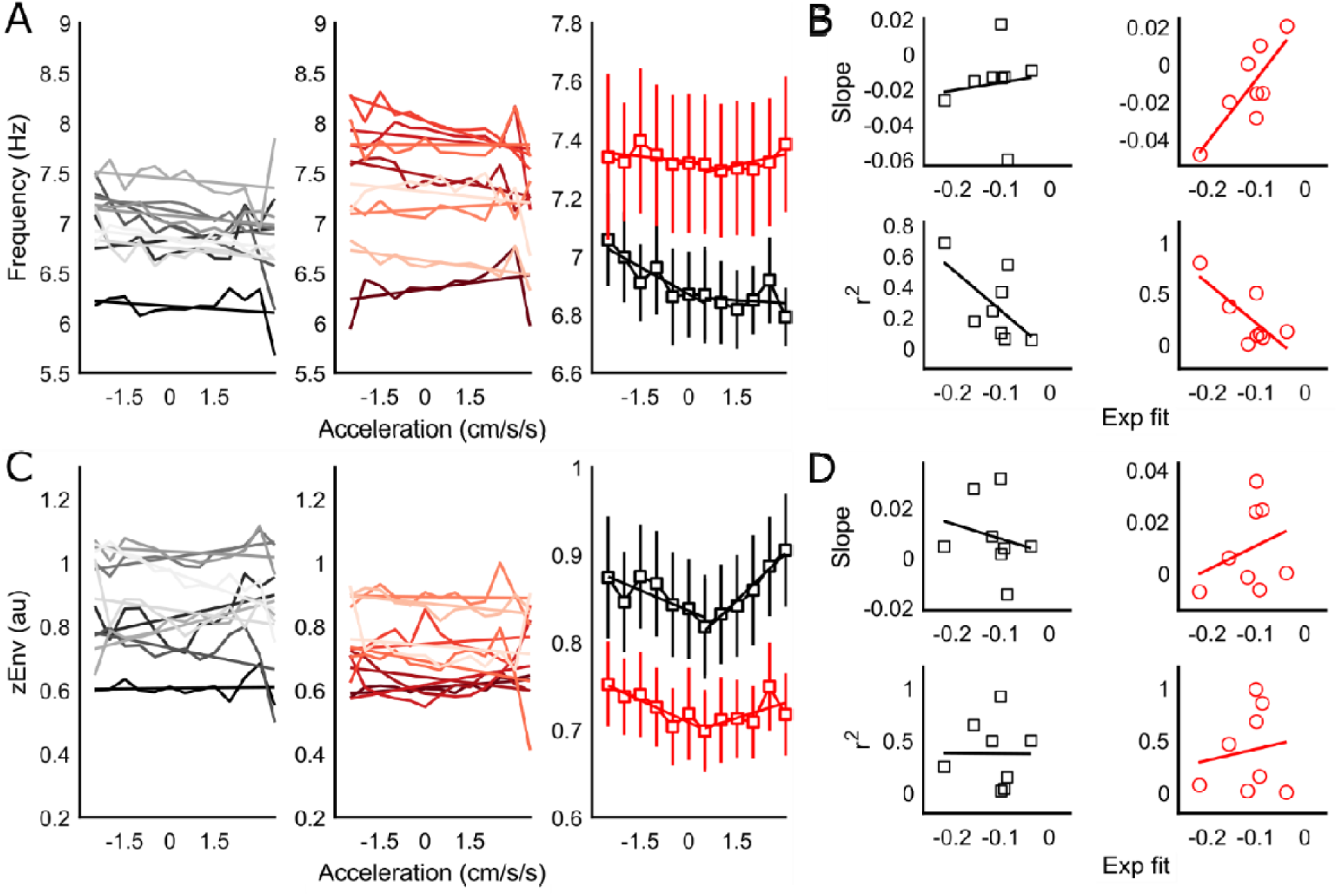
Acceleration modulation of hippocampal and supramammillary theta oscillations. **A**: Raw data and their linear fits for the hippocampus (right, black) and the supramammillary area (middle, red) theta frequency across binned acceleration. Population data and their fits are depicted in the right panel. **B**. Correlations between the magnitude (slope, top) and robustness (*r*^*2*^, bottom) of the frequency versus acceleration scaling and water maze learning (exponential fit slope parameter) for hippocampal (black, left) and supramammillary (red, right) theta. **C**: Same as in **A** but for acceleration versus theta power correlations. **D**: Same as in **B** but for theta power versus acceleration scaling and water maze performance correlations.

Both the HPC (Figure 5C, right) and SuM (Figure 5C, middle) show weak and variable associations between theta power and acceleration. At the population level, the data suggest a clear quadratic relationship between acceleration and theta power (Figure 5C, right). This observation is supported by much stronger association between negative (*r*^*2*^ = 62, F(10) = 8.09, *p* = .04) or positive acceleration (*r*^*2*^ = 98, F(10) = 165.93, *p* < .001) with HPC theta power alone, rather than a linear fit using the overall acceleration (*r*^*2*^ < .001, F(10) = .09, *p* = .76). Although a similar relationship can be observed for SuM theta power and acceleration, the negative acceleration (*r*^*2*^ = .85, F(10) = 29.33, *p* = .003) yielded a better fit than the positive (*r*^*2*^ = 40, F(10) = 2.62, *p* = .18), as well as the overall acceleration (*r*^*2*^ = .06, F(10) = .67, *p* = .43). Given the relationships between acceleration and theta power are clearly not linear and appear to be quadratic, we performed linear fits on the absolute values of acceleration and extracted the slope and *r*^*2*^ to regress against water maze learning estimate. Similar to HPC theta power relationships with speed, neither slope (*r*^*2*^ < .05, F(10) = .30, *p* = .60) nor *r*^*2*^ (*r*^*2*^ < .001, F(10) < .001, *p* = .99) of absolute acceleration fits correlated well with the rate of spatial learning (Figure 5D, right panels). The same is for the SuM, showing no correspondence between the slopes (*r*^*2*^ = .09, F(10) = .60, *p* = .47) and *r*^*2*^ (*r*^*2*^ = .02, F(10) = .13, *p* = .24) extracted from absolute acceleration and water maze learning (Figure 5D, right panels). A summary between linear scaling of theta variables and their relationship with water maze learning are presented in Table 1.

### Contributions of spatial and learning metrics to ongoing theta frequency and power

From our regression analyses, it appears both theta variables have better correlates with speed rather than acceleration, and that how HPC theta frequency scaling with swimming speed serve as a superior predictor for water maze performance compared to other possible combinations. As an alternative way of characterising a potential link between speed modulation on HPC theta frequency relating to learning, as well as circumventing the need to perform multi-step regressions with dimensionally reduced data (i.e. regressing fit parameters against another), we performed GLM with binned speed, binned acceleration and our novel performance metric as potential predictors for theta measures for each rat. For theta frequency versus swimming speed (Figure 6A), zβ were all positive for HPC except for one rat. As a group, the zβ were significantly different to zero (t(7) = 3.05, *p* = .02). This was not the case for SuM theta frequency (t(7) = −.04, *p* = .68). Acceleration was a poor predictor for both HPC (t(7) = 96, *p* = .38) and SuM (t(7) = −.53, *p* = .61) theta frequency. Interestingly, the water maze performance metric appears to be the best predictor for theta frequency in both the HPC (t(7) = 5.85, *p* < .001) and the SuM (t(7) = 2.59, *p* = .04).

**Figure 6.**
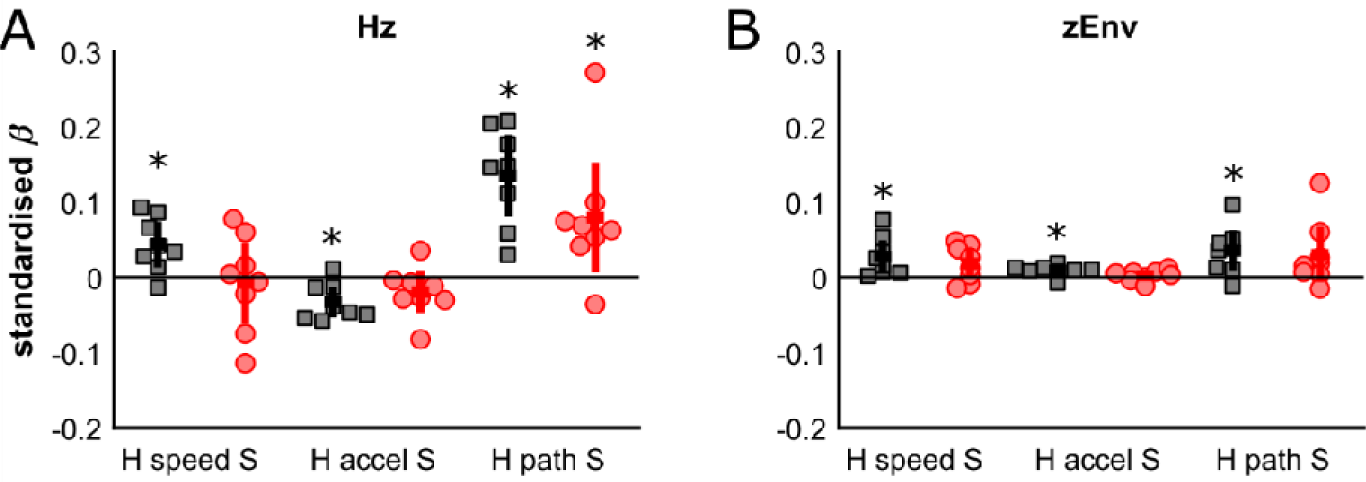
Standardised betas from GLM. **A**: Betas from each predictor variable from each brain structure for theta frequency. **B**. Same as **A** but for theta power. H: hippocampus; S: supramammillary area. Asterix denotes *p* < .05 with Student’s t-tests.

For the same analyses relating to theta power, speed appeared to be a good predictor for both HPC (t(7) = 2.99, *p* = .02) and SuM (t(7) = 2.20, *p* = .06) theta power (Figure 6B). Acceleration is the best predictor for HPC theta power (t(7) = 3.59, *p* = .009) but not for the SuM (t(7) = 1.01, *p* = .35). Water maze performance also served as a good predictor for HPC theta power (t(7) = 3.08, *p* = .02), but less so for the SuM (t(7) = 1.96, *p* = .09).

## Discussion

Much theory and empirical data suggest HPC theta oscillations, particularly concerning locomotion speed, have a role in spatial navigation (Montgomery et al., 2009; Burgess and O’Keefe, 2011; Richard et al., 2013; Schmidt et al., 2013; Belchior et al., 2014; Hernandez-Perez et al., 2016; Korotkova et al., 2018). We used the rate of water maze learning as a functional test for spatial navigation. We were able to show: 1) HPC theta frequency and power scale linearly with swimming speed; 2) HPC theta frequency and power have opposite spatial distributions in the water maze; 3) more robust HPC theta frequency scaling with swimming speed is correlated to better water maze learning; 4) swimming speed and goal-directed spatial navigation are the best predictors for HPC theta frequency; and, 5) swimming speed, acceleration and goal-directed spatial navigation all contribute to HPC theta power and frequency predictions. Our findings are consistent with the notion that HPC theta reflects inherent path integration abilities in individuals, which predict the rate of water maze learning.

We were able to show that swimming speed correlates to, and is a useful predictor of, HPC theta frequency. Previous studies have reported a lack of robust linear correlation between HPC theta frequency and swimming speed (Whishaw and Vanderwolf, 1971; Hernandez-Perez et al., 2016). However, the conclusions were either made qualitatively (Whishaw and Vanderwolf, 1971) or based on a longer 2 s window spectral estimate of theta frequency that was combined with an unknown undescribed treatment of tracking data at a higher rate (Hernandez-Perez et al., 2016). Since we employed “instantaneous” theta frequency and power derived from the analytic signal in our study, the increase in precision in the temporal domain may have unmasked inherent speed modulation of HPC theta frequency that exists in swimming animals. However, it is clear from converging studies that uncoupling movement speed from motor output results in the loss/flattening of the HPC theta frequency versus speed relationship (Czurko et al., 1999; Shin and Talnov, 2001; Terrazas et al., 2005; Kuo et al., 2011; Ravassard et al., 2013); it remains to be seen if movements at the same speeds between swimming and running animals would differ in the strength (i.e. slope) of the scaling function.

Our data indicate movement speed does partially account for variance in HPC theta frequency, but estimates of water maze learning appear to provide more predictive value. This is in line with previous studies, where GLM identified movement speed and location on the maze, but not acceleration, as robust predictors of HPC theta frequency and to a lesser extent, power, in spatial tasks (Montgomery et al., 2009; Schmidt et al., 2013). A different study examining HPC theta frequency versus movement speed relationships also revealed that the slope and *r*^*2*^ were both correlated with spatial alternation performance (Richard et al., 2013). More indirectly, when movement speeds are matched, there is an apparent increase in HPC theta power and frequency correlating to correct spatial choices in a foraging task (Belchior et al., 2014). Our findings reported here are consistent with the idea that HPC-dependent task performance metrics are highly correlated with HPC theta frequency but less so with power. HPC theta frequency is known to be less variable than power across the whole hippocampus (Maurer et al., 2005; Montgomery et al., 2009; Hinman et al., 2011), hence it is not surprising measures of kinetics and spatial learning correlated better with HPC theta frequency than power.

In our study, the behavioural context is one of spatial learning. It has been documented previously that exposure to novel spatial environments reduces the slope of the HPC theta frequency versus movement speed relationship (Jeewajee et al., 2007; Wells et al., 2013). Unfortunately, limitations imposed by our study design prevent a conclusive demonstration of increases in HPC theta frequency versus speed slopes as a function of learning (i.e. progression from “novel” to “familiar” environments). Due to the nature of the water maze learning task, less data become available as animals learn to successfully locate the hidden platform. We were able to detect a correlation between the slope of HPC theta frequency versus speed and path length (data not shown). However, high variances in the direction (i.e. positive/negative slopes) and value of fitted slope as a function of path length, presumably driven by the paucity of data in shorter trials, prevented any useful conclusions to be drawn. Future studies utilising probe trials at the end of training would be useful in providing adequate data to investigate if increases in slope of the HPC theta frequency versus swimming speed relationship emerges as animals acquire the position of the hidden platform (i.e. as the novel spatial environment becomes familiar). Relocation of the hidden platform would also be useful in disambiguating effects of spatial novelty on HPC theta frequency versus speed scaling and learning a new goal location.

We have demonstrated that the HPC theta frequency versus swimming speed relationship can predict learning rates in the water maze. It is currently unclear if the predictive value is restricted to swimming speeds, or whether it can be generalised to the strength of HPC theta frequency versus movement speed on dry land. A crucial experiment validating our proposed importance of HPC theta frequency versus speed scaling in spatial cognition would be to examine if the strength of frequency versus speed coupling on land is predictive of water maze learning rate in individual animals. If so, it would suggest the slope and to a certain extent, measures of goodness-of-fit (e.g. *r*^*2*^) of HPC theta frequency versus speed relationship may serve as a potential biomarker for spatial cognition and its deficits. In addition, the use of a multi-day instead of a single-day paradigm used in the current study would also provide further insight on if the relationships described here is generalisable in water maze learning and goal-directed spatial navigation and not dependent on unknown features of the single-day implementation.

A potential confound in our study is the effect of body temperature reduction in our single-day, 16 consecutive trials version of the water maze. Previous studies have shown a reduction in body temperature reduces HPC theta frequency (Whishaw and Vanderwolf, 1971; Wells et al., 2013). We have previously demonstrated that the effect of cooling on HPC theta frequency follows a strong exponential decay function (Pan and McNaughton, 1997), which is also the type of curve fitting we performed to assess learning rates in the current study. Given we did not monitor the rats’ body temperature, how changes in body temperature throughout our water maze training impact on our conclusions cannot be accounted for. However, we note that our GLM analysis is not directly impacted by the exponential decrease in path length and body temperature as a function of time. With our novel real-time water maze performance metric, the GLM analysis suggested that our instantaneous performance metric can still predict changes in HPC theta frequency, independent of shared similarities between path length and body temperature time-courses.

Although the HPC theta frequency versus movement speed relationship is well-described, it is not the only speed representation available to the canonical spatial circuits. Speed information is also represented in firing rates (Kropff et al., 2015; Gois and Tort, 2018) and burst frequency (King et al., 1998) of single cells along the spatial circuit the HPC is a part of, and appear to coexist as a competing source of speed information in the medial entorhinal cortex (Hinman et al., 2016). Some experimental evidence suggests self-motion- and sensory-derived (mostly visual) signals may underlie the existence of multiple speed information in the spatial circuit (Campbell et al., 2018; Jayakumar et al., 2019). The nature of the speed information that the CA1 theta LFP carries is unclear; however, this theta appears to be essential for spatial representation in the HPC and medial entorhinal cortex (Brandon et al., 2011; Koenig et al., 2011; Bolding et al., 2019), and uncoupling of self-motion and actual movement appear to suppress its relationship with speed (Czurko et al., 1999; Shin and Talnov, 2001; Terrazas et al., 2005; Kuo et al., 2011; Ravassard et al., 2013). Of interest, there is no reported change in place cell firing rates in the water maze in relation to speed (Hollup et al., 2001a, b). Given local field potentials are epiphenomena and a composite surrogate measure for local excitability, it is clear from the current and other studies that HPC theta, particularly frequency, carries speed-related information among other types. Additional experiments are needed to tell if and how multiple speed codes in the HPC represent different speed representations such as self-motion, perceived movement through space, and physical effort related to movements.

Human HPC theta oscillations are also correlated to movement through space (Ekstrom et al., 2005; Bush et al., 2017), mnemonic processes (Rutishauser et al., 2010) and spatial navigation (Bush et al., 2017). Importantly, the removal of salient sensory cues in virtual reality environments lowers the frequency of human hippocampal theta (Bohbot et al., 2017), which may reflect a decrease of the theta-speed relationship as seen in rodents. Overall, there is a good correspondence between human and rodent data on how HPC theta relates to movement kinematics (Kunz et al., 2019). There is also evidence that such relationships can be exploited in clinical settings to assess the effectiveness of anxiolytics in the HPC (Wells et al., 2013; Young and McNaughton, 2020) and beyond. An improved understanding of how the HPC theta frequency versus speed relationship is generated, modulated and utilised in the brain can provide fundamental insights to sensorimotor integration (Bland and Oddie, 2001) in health and disease.

### Conclusions

Previous studies have shown behavioural tasks can significantly modulate HPC theta oscillations, beyond its correlations with movement speed. Our findings take a step further in providing evidence to support the idea that HPC theta frequency correlations with speed may be actively utilised to support spatial cognition. This appears to be a specific effect as: 1) such relationship was weak or inconsistent between different analytical approaches for HPC theta power; 2) acceleration cannot be used for reliable HPC theta frequency and power predictions and; 3) identical analyses carried out on SuM theta provided no evidence for kinematic scaling and their relationship to water maze learning. Thus, we conclude that HPC theta frequency and speed correlations serve its theorised role (Burgess and O’Keefe, 2011) in providing a distance metric in path integration for spatial cognition.

## Acknowledgments

This work was supported by the Marsden Fund of New Zealand (UOO105) and New Zealand Neurological Foundation (0024/PG/McNaughton).

## Contributions

CKY conceived the study, carried out the analyses and part of the histology. MR collected the behavioural, neurophysiological and histological data. CKY and NM prepared the manuscript.

